# Distributed network sensing as an escape strategy

**DOI:** 10.64898/2026.11.05.764324

**Authors:** Mathieu Le Verge-Serandour, Alexandra Bienau, Friedrich C. Simmel, Karen Alim

## Abstract

The response of a swarm of ants or birds, living tissue of cells, or a robotic swarm, to spatial signals from the environment as a whole is a fundamental challenge for these decentralized systems. A promising alternative is embodied distributed sensing and decision-making, so that local environmental cues directly shape coordinated reconfiguration, growth, or movement without centralized control. Fluidic networks offer a physical basis for such systems, yet existing implementations are predesigned and limited in their ability to process spatial information across extended systems. Here, we investigate how fluid transport networks enable environmental sensing and adaptive response using the slime mould *Physarum polycephalum*. By exposing *Physarum* vein networks to dynamic pH environments, we show that the organism’s ability to respond to aversive stimuli is governed by its spatial extent and network connectivity. Above a certain size, organisms with network morphology reliably detect and escape harmful conditions by network reconfiguration, whereas disconnected or spatially confined organisms fail to respond. In environments with competing stimuli, network morphologies enable integration of spatial information and repositioning into favorable regions. We identify local contractions as a physical mechanism that drives fluid redistribution and converts local stimulation into system-level responses. Finally, we generate different morphologies using fluidic and electrochemical control, demonstrating large-scale adaptation to complex environments. These results establish distributed fluidic networks as a framework for integrated sensing and actuation, providing design principles for autonomous soft robotic systems in dynamic environments.

**Significance:** To sense across a large area, a living system either builds a sensor everywhere, at high cost, or grows a sparse network that samples space cheaply. The slime mould *Physarum polycephalum* takes the network route, and unlike most adaptive materials, it rebuilds its entire architecture within minutes, not hours or days. We show that this network architecture is exactly what enables *Physarum* to quickly detect and escape a localized chemical threat. We identify the physical mechanism driving escape: tubes near the threat contract, raising local pressure and pushing fluid mass away from danger and into the rest of the network, with no central controller directing the response.

---

Decentralized systems, from swarms of ants and birds to living tissues and robotic collectives, must often integrate signals distributed across space into a coordinated, system-level response without relying on a central controller or authority. Achieving this robustly under dynamic, unstructured conditions remains a major challenge despite substantial advances in locomotion and control. Such environments require systems not only to detect spatially distributed stimuli but also to integrate this information and translate it into coordinated, adaptive responses. Multi-agent systems and robotic swarms achieve robustness through distributed coordination among discrete agents, whereas soft robots exploit compliant materials and embodied mechanics to interact adaptively with uncertain environments. However, integrating spatially distributed environmental signals into coordinated physical adaptation across an extended body or collective remains difficult. Identifying their underlying principles of such distributed adaptation, spanning not only movement, but also morphology, spatial extent, and internal organization in response to changing conditions, remains an open problem to living and synthetic systems.

## Fluidics for sensing, actuation, and computation

Fluidics has emerged as one of the key modalities for sensing, actuation, and control in soft and compliant systems (1, 2). Pressures and flows can be directly coupled into compliant bodies, enabling distributed actuation and large deformations (3, 4). Building on this principle, fluidic systems have demonstrated capabilities such as flow-based logic and signal processing (5, 6), as well as fully soft autonomous operation, exemplified by the octobot, in which microfluidic circuits regulate fluid flow without electronics (7). More recently, fluidic sensing strategies have shown that environmental interactions can be inferred from pressure responses without embedded sensors (8).

Despite these advances, current fluidic implementations remain largely predesigned, static architectures and are typically limited to local sensing. While they enable actuation, timing, and logic, they do not yet provide mechanisms for distributed environmental sensing and decentralized information processing across spatially extended systems. As a result, strategies that couple transport, sensing, and adaptive response in a fully integrated manner remain largely unexplored.

## Biological transport networks

In contrast, many biological systems routinely achieve robust and adaptive behavior in complex environments through distributed transport networks composed of branched, interconnected veins. These network architectures are widespread across organisms, from fungal mycelia and plant vasculature to circulatory systems, and support not only the transport of nutrients and waste but also long-range communication through fluid flow. Signals carried by transported molecules, pressure variations, or chemical gradients propagate across the network, effectively coupling spatially separated sensing sites into a coordinated system (9). This enables organisms to detect spatial variations and reorganize their morphology to maintain function (10). Fluid transport may be driven by centralized pumps, as in the cardiovascular system, or by decentralized pumping or oscillatory transport mechanisms, as observed in pancreas (11), plants (12), fungi (13), and slime molds (14), allowing coordination across large spatial scales analogous to communication between distributed agents.

A defining feature of biological transport networks is their ability to grow and remodel. Unlike fixed engineered infrastructures, living vascular systems continuously modify the size, connectivity, and number of veins in response to internal demands and environmental constraints. Such restructuring enables exploration–exploitation strategies, where networks expand into favorable regions while retracting from unfavorable ones (15). These adaptive processes follow optimization principles that balance transport efficiency, robustness, and energetic cost (16–18). In parallel, transport networks maintain stable internal conditions by regulating internal composition and flow distribution, coupled to exchanges with the environment. Local imbalances in flow, composition, or transport demand may act as system-wide signals that trigger structural adaptation, linking environmental sensing directly to physical reorganization (19, 20).

The importance of vascular transport has been highlighted as a major challenge for scaling functional engineered tissues and adaptive materials (21). However, despite the ubiquity of transport networks in biology, the principles by which system size and network architecture enable distributed sensing and coordinated response remain poorly understood.

## Our approach: *Physarum*

Here, we investigate a living vascular network in dynamic chemical environments and extract strategies for integrating distributed stimuli information into network morphology. We use the unicellular slime mould *Physarum polycephalum*, which forms dynamic, reconfigurable transport networks spanning sizes from hundreds of micrometers to several meters.

Despite lacking a nervous system, *Physarum* exhibits a wide range of adaptive behaviors, including distributed decision-making during foraging (22, 23), nutrient optimization (24, 25), and the ability to solve spatial optimization problems through network reconfiguration (26, 27). It can furthermore display primitive cognitive behaviors such as habituation and learning (28–31). *Physarum* responds to a wide range of environmental stimuli, including light, nutrients, and chemical signals (32), and dynamically adapts its transport morphology to complex environments. Internally, rhythmic contractions drive fluid transport and signal propagation throughout the network (33, 34), and the resulting network morphology can encode spatial information about past environmental conditions (35, 36). This tight coupling between flow, structure, and environment makes *Physarum* a unique and powerful model platform to study distributed sensing in physical networks in the absence of centralized control.

## This work: pH environments

In this work, we expose *Physarum* to spatially structured and dynamically evolving pH fields. *Physarum* preferentially grows within a narrow pH range (pH 4.5-7) and avoids conditions outside this range, providing a well-defined stimulus for probing environmental sensing.

We find the organism to effectively escape regions affected by strongly acidic pH while growth is promoted in areas exhibiting the preferred pH range (Fig. 1C). We further demonstrate that the ability to respond to an aversive stimulus is governed by spatial extent and network connectivity (Fig. 1E). *Physarum* networks can furthermore resolve competing stimuli in complex, spatially organized environments, and we demonstrate adaptation and growth into predicted corresponding morphologies (Fig. 2, Fig. 4). By analyzing the underlying dynamics, we identify a physical mechanism in which local contractions respond to pH and drive fluid redistribution (Fig. 3), enabling coordinated avoidance and adaptation across *Physarum* networks. Even in dynamically evolving environments, *Physarum* networks continuously adapt to changing spatial patterns (Fig. 5).

**Fig. 1.**
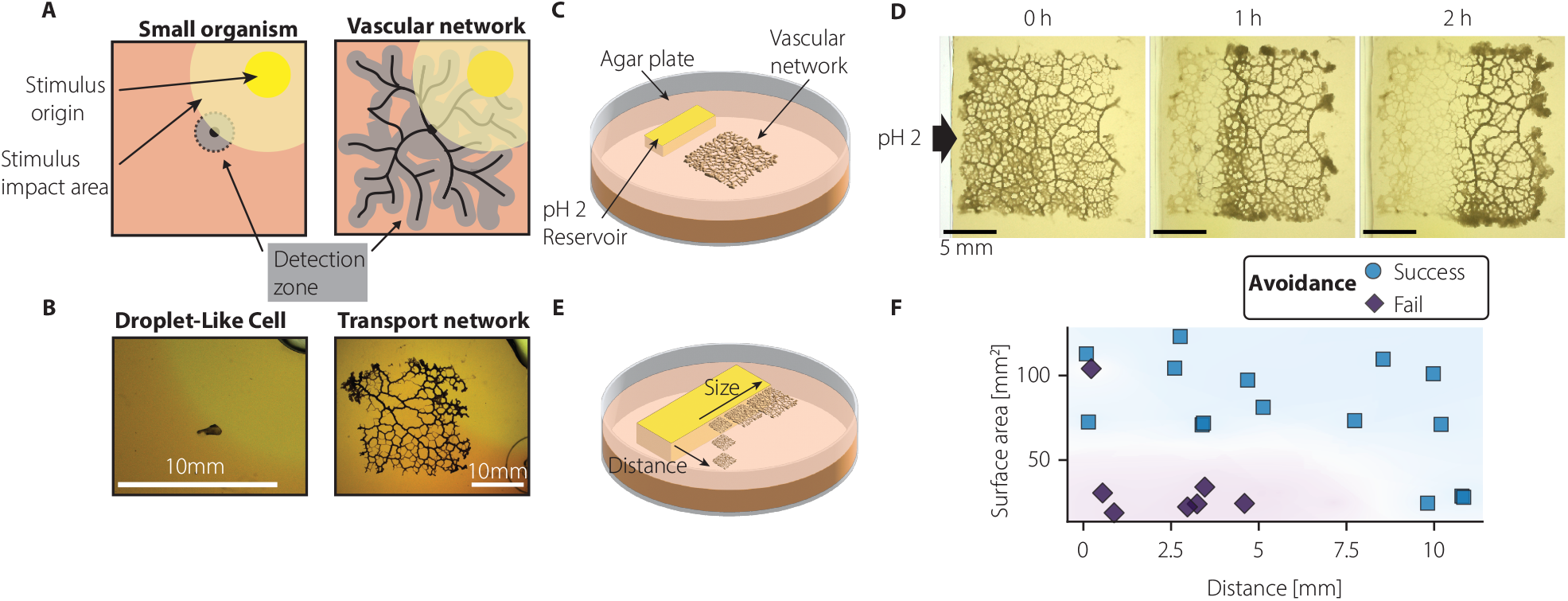
*Physarum* network size enables escape from harmful pH gradients. (**A**) While a small organism is limited by the diffusion of aversive stimuli, *Physarum* networks can efficiently detect it. (**B**) Examples of *Physarum polycephalum* at different scales, with (left) droplet-like cell and (right) transport network. (**C**) Avoidance assay: a pH 2 reservoir carved into agar generates a diffusive acid gradient advancing to the network. (**D**) Representative escape of a network to an advancing acid front. Scale bar, 5 mm. (**E**) Avoidance assay showing variation in organism size and distance to the source. (**F**) Escape of networks as a function of network’s surface area and distance to pH reservoir reveals that size limits success in escape. Shaded regions are guides to the eye.

**Fig. 2.**
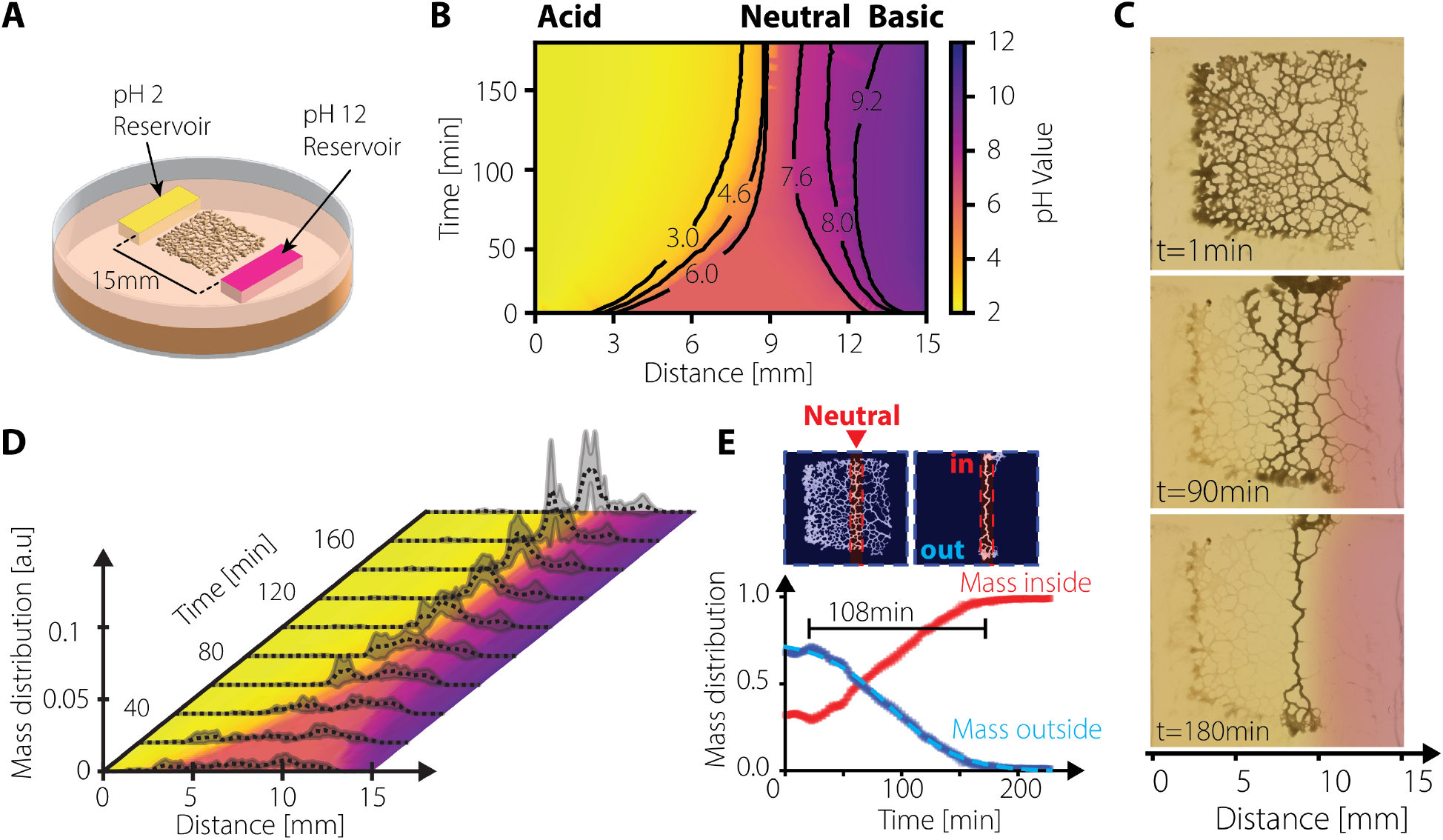
Network reorganization enables navigation in opposing pH gradients. (**A**) Experimental setup with opposing reservoirs at pH 2 and pH 12 surrounding a *Physarum* network. (**B**) Reconstructed kymograph of pH evolution, showing converging acid (pH<6) and basic fronts (pH>8) that define a central neutral corridor. (**C**) Representative network reorganization, with the network aligning along the neutral region. (**D**) Temporal evolution of biomass distribution (N = 4) overlaid on the pH map, showing progressive localization within the neutral corridor. (**E**) Fraction of mass inside and outside a predefined region of neutral pH, illustrating avoidance of extreme pH conditions over 180 min.

**Fig. 3.**
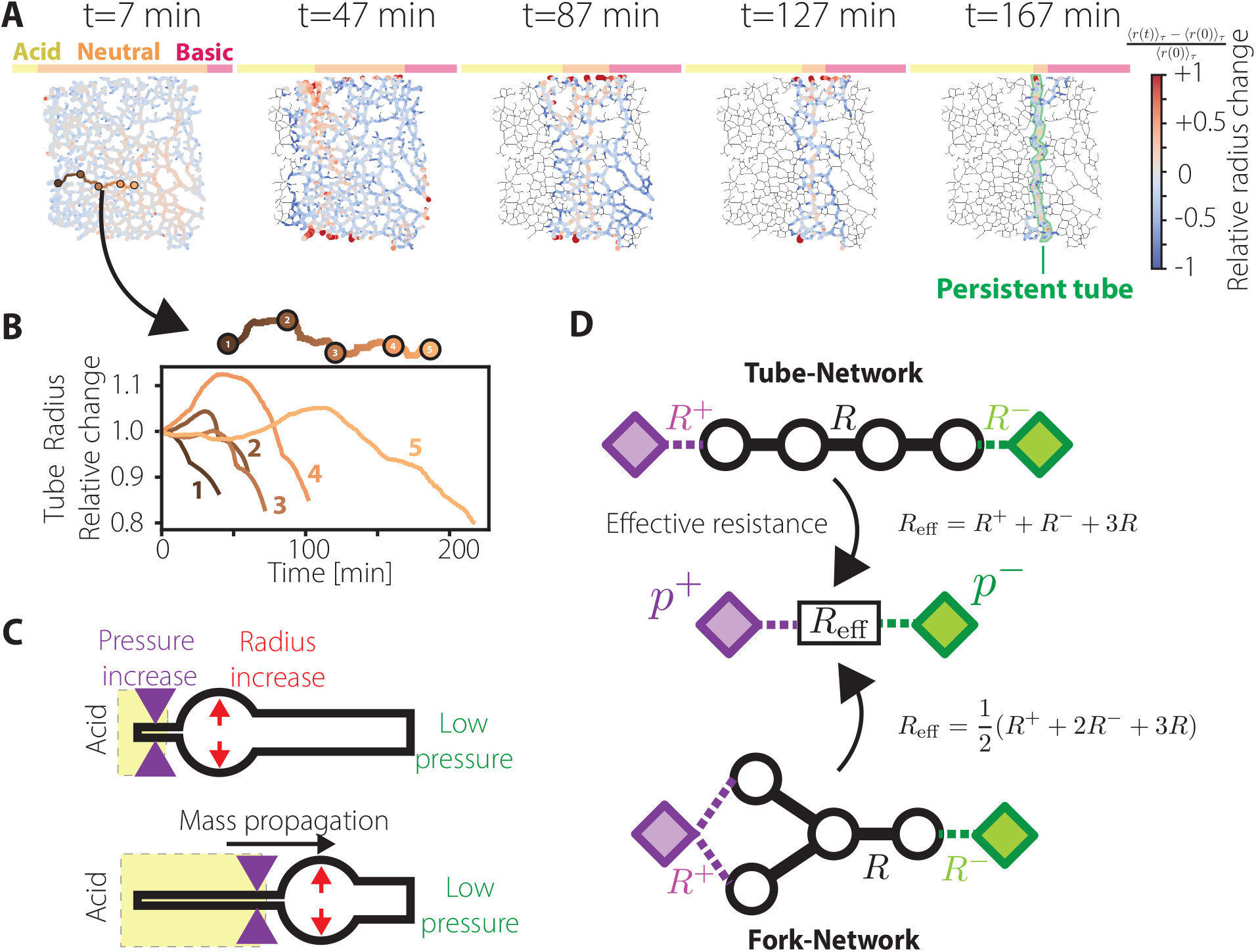
Avoidance drives adaptive reorganization of transport networks. (**A**) Networks escape unfavorable pH (acid: yellow, pink: acid) by changing its veins’ radius over time. Color encodes averaged (over 4 min) relative radius normalized by the averaged initial radius. (**B**) Selecting five positions along a path from acid-exposed veins to the left toward persistent veins located in the favorable pH zone, the vein radii show a rightward propagating local increase before shrinkage, (**C**) which can be schematized as a pulse-like propagation of biomass from high pressure, arising from stimulus-induced vein contraction, to low pressure in unaffected veins. (**D**) Extending to elementary network architectures, a tube and a fork, connected to pressure sinks (green, *p*^−^) and sources (purple, *p*^+^), shows that the effective resistance of the fork is only half of the tube, facilitating a more efficient escape for forked network architectures over purely linear veins.

**Fig. 4.**
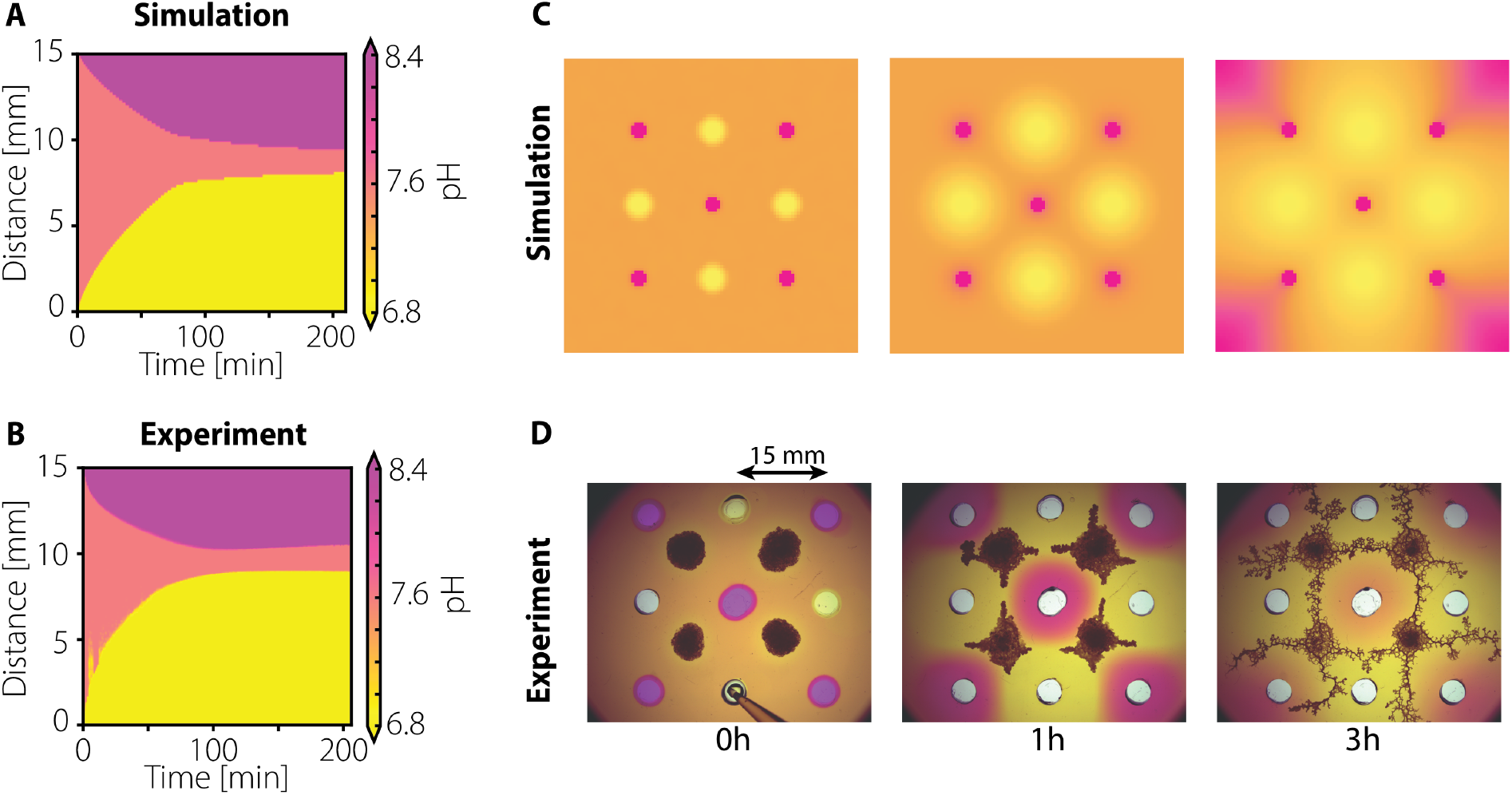
Fluidic control of complex pH landscapes. (**A**) Simulated and (**B**) experimental kymographs of acid–base diffusion from opposing fluidic sources show close qualitative agreement. (**C**) Multiple reservoirs carved into agar generate complex pH geometries, which guide (**D**) the formation and organization of *Physarum* vein networks.

**Fig. 5.**
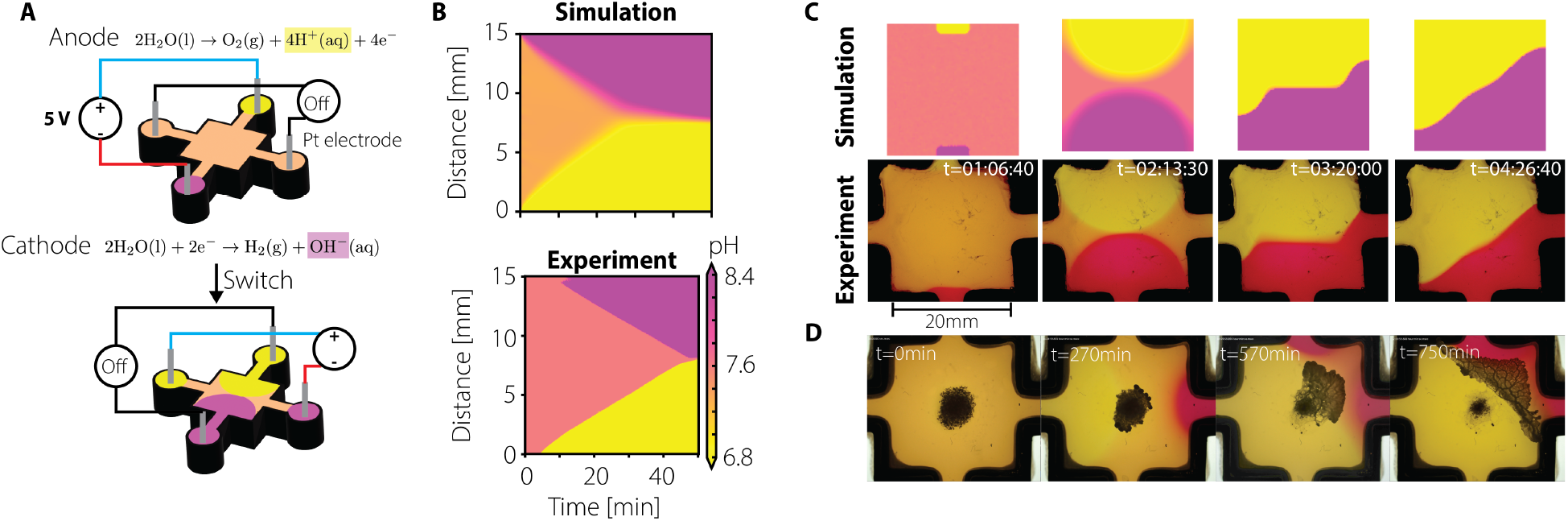
Electrochemical control of dynamic pH landscapes controls network dynamics. (**A**) Electrochemical control of dynamic pH landscapes is implemented using a 3D-printed chamber with independently addressable platinum electrodes. (**B**) Simulated kymographs of the electro-diffusive system reproduce the experimentally observed dynamics and guide experiment design. (**C**) Sequential activation of orthogonal electrode pairs enables dynamic reconfiguration of pH patterns, shown in simulations (top) and experiments (bottom). (**D**) Dynamic control of *Physarum* vein network: droplet-like cells fuse, reorganize and extend protrusions along electro-diffusively defined neutral regions as the imposed field (DC 3 V) is reoriented in time.

With these results, we establish distributed fluidic networks as a physical framework for integrating spatial sensing and actuation, providing design principles for autonomous soft robotic systems capable of operating in complex environments.

## Results

### Size matters: distributed sensing enables escape from harmful environments

Timely detection of environmental signals and appropriate responses are essential for biological and robotic systems alike. At the molecular level, this ability is fundamentally constrained by diffusion, which limits how quickly and how far chemical signals propagate through space. As a result, spatially limited systems may respond too late to environmental cues. One potential solution is to distribute sensing across space: network-like architectures extend the effective detection range by sampling the environment at multiple locations simultaneously (Fig. 1A).

We use *Physarum polycephalum*, a unicellular organism capable of spanning sizes from hundreds of micrometers to several meters. Depending on its life cycle, it exists either as small, droplet-like cells, or as extended, branched transport networks. We exposed this system to diffusion-generated pH gradients to examine how size and morphology shape environmental signal detection and response. *Physarum* preferentially grows within a narrow pH range (pH 4.5-7) and avoids aversive conditions outside this range (37, 38) providing a well-defined stimulus for probing environmental sensing. In our setup, localized pH 2 reservoirs embedded in agar (Fig. 1B) generate pH fronts that advance toward the samples by diffusion. Upon exposure, *Physarum* networks rapidly evacuate their biomass from aversive regions and establish new expansion regions at distant locations (Fig. 1C).

We exposed trimmed *Physarum* networks spanning sizes from approximately 20 to 125 mm^2^ to diffusive acid gradients, revealing a sharp size-dependent transition in behavior (Fig. 1D-E). *Physarum* networks with areas larger than approximately 70 mm^2^ reliably detected the approaching acid front and retreated, whereas smaller *Physarum* networks failed to respond. Increasing the distance to the source reduced the strength of the pH stimulus, enabling smaller networks to escape. Beyond approximately 7.5 mm, all samples avoided the advancing front before the stimulus induced sample inactivity.

To disentangle the roles of size and connectivity, we examined droplet-like cells of *Physarum* for comparison. Individual droplet-like cells typically have sizes of 10 mm^2^ or lower and largely failed to respond, entering an inactive state upon exposure. Using the ability of *Physarum* droplets to fuse into larger droplet-like cells, with size comparable to larger *Physarum* networks, we did not observe a significant improvement in avoiding harmful conditions, indicating that surface coverage alone is insufficient for reliable response (Fig. S3).

### Navigation and redistribution in opposing pH gradients

To probe how networked systems adapt to competing stimuli, we introduced *Physarum* networks into environments containing two opposing chemical cues: reservoirs buffered at pH 2 and pH 12 (Fig. 2A). The agar and reservoirs were supplemented with the pH indicator phenol red, which turns pink under basic conditions (pH > 8.4). Initially, at homogeneous pH (pH 6.4-8.4), the conditions in the agar evolved as unfavorable pH fronts propagated from their respective reservoirs. Between the two reservoirs (separated by 15 mm), these fronts converged to form a narrow corridor of near-neutral pH, flanked by regions of extremely unfavorable pH (Fig. 2B, Movie S1).

We quantitatively reconstructed the spatiotemporal evolution of this pH landscape using four complementary indicator dyes that together cover the pH range from 3.0 to 9.2 (Fig. S4, Movies S2-S5). Interpolation of these measurements yielded a continuous pH map, from which we extracted effective diffusion coefficients of acid *D*_*a*_ = 9 × 10^−5^ cm^2^/s and base *D*_*b*_ = 5 ×10^−5^ cm^2^/s, consistent with literature values (39), see Fig. 2B.

When trimmed *Physarum* networks, standardized to nearly 1 cm^2^ (39), were placed at the center of this environment, 2.5 mm from each reservoir, they dynamically reorganized in response to the evolving gradients. Initially isotropic, the networks progressively elongated and aligned along the neutral corridor defined by the opposing fronts (Fig. 2C).

To quantify this behavior, we measured the spatiotemporal distribution of biomass across multiple samples and compared it to the reconstructed pH field (Fig. 2D). While mass was initially broadly distributed, it rapidly concentrated within a narrow region corresponding to pH 4.05 to 7.26 (Fig. S4E). The escape dynamics, measured by the fraction of mass within this region over time, revealed a characteristic timescale of approximately 108 min, with total mass localized within the neutral corridor (Fig. 2E). This timescale is comparable to both previously reported *Physarum* responses to aversive stimuli (40, 41) and the diffusion timescale of H_3_O^+^ and hydroxide (OH^−^) ions (approximately 160 min), indicating that network reorganization is tightly coupled to environmental dynamics.

### *Physarum* network topology offers best chances for efficient avoidance

While *Physarum* networks efficiently reposition in complex environments by tracking favorable pH regions, this adaptation necessarily relies on rapid internal reorganization. As the pH landscape evolves, the *Physarum* network must continuously redistribute mass and reshape its architecture to maintain function and avoid harmful regions. We therefore examine the dynamics of this response by quantifying how vein radii evolve during avoidance.

Mass redistribution emerges from asymmetric contractions that redirect flow from exposed regions toward persistent veins via evacuating pathways. In *Physarum*, peristaltic contractions generate internal flows that drive mass transport (33). Over longer timescales, shear-stress-dependent adaptation leads to the pruning of veins with low flow (42). Upon exposure to an external stimulus, contraction frequency and amplitude increase noticeably, reflecting an active response (32). To quantify these response dynamics in our test network, we segmented individual vein cross-sections and tracked their radii over time, using the bright-field intensity as a proxy for vein radius, see Fig. S5B & C (41). We identified one central persistent vein in the neutral region, which collected the evacuating mass, while all other veins were removed as unfavorable pH diffused (Fig. 3A).

Focusing on veins connecting the leftmost periphery to the persistent veins in near-neutral pH, we analyzed the relative radius change at several positions along this path. As shown in Fig. 3B, the radii exhibit a pronounced increase followed by a gradual decay along the path, consistent with a travelling wave of mass propagating from the left to the persistent vein. Repeating this analysis for other parallel paths reveals a similar trend for both unfavorable pH conditions when combined with contraction analysis, although the response along the basic path was delayed relative to the stimulus onset (Fig. S5D-F).

These observations are consistent with a local pressure increase generated in response to the advancing acid front (Fig. 3C). Exposure to the advancing aversive stimulus causes veins, that sense the unfavorable pH, to contract, thereby increasing local pressure. This pressure buildup drives mass redistribution out of contracting veins and ensues dilation of neighboring veins until neighboring veins sense the unfavorable pH and, themselves contract and locally increase pressure. The observed transient increase in vein radius amplifies the pressure difference and, thus, acts as an actuation mechanism. The resulting dynamics resemble a pulse-like propagation: as the stimulus diffuses underneath the *Physarum* network, tube contraction and forward dilation progress toward the persistent veins in the near-neutral region.

Finally, we note that branched network architectures are inherently more efficient for such escape dynamics. We compared two elementary graph topologies with identical volumes, defined by the same number of veins with equal hydraulic resistance (Fig. 3D). By analytically computing the effective resistance between fixed boundary pressures, with the higher-pressure *p*^+^ representing an aversive-exposed vein and the lower pressure *p*^−^ representing a persistent vein, we find that a fork-network is 66% more efficient than a linear network, see *SI Appendix* for details.

### Quantitative evolution of diffusive pH through fluidic control

While *Physarum* networks can robustly adapt to static or slowly evolving environments, real-world conditions are often dynamic and require continuous adjustment. We therefore seek to impose pH landscapes that can be actively changed over time, enabling the system to probe whether *Physarum* can autonomously track and respond to continuously evolving stimuli.

To better control and predict the emergence of pH patterns, we model the coupled transport of acid and base using a reaction-diffusion framework implemented in a dedicated Python solver. The system is defined on a two-dimensional square domain of size *L*, with acid concentration *a*(*x, y, t*) and base concentration *b*(*x, y, t*) obeying

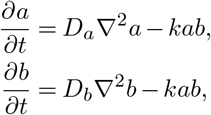

where *D*_*a*_ and *D*_*b*_ are the diffusion coefficients of the acid and base, respectively, and *k* is the rate of the neutralization reaction. This formulation corresponds to a two-species annihilation process *A* + *B* → ∅ (43). An acid influx *J*_*A*_ is imposed on the left boundary, and an equal base influx *J*_*B*_ on the right boundary, such that *J* = *J*_*A*_ = *J*_*B*_ = 10^−9^ mol/(cm^2^ s), ensuring a symmetric configuration. We first reproduce the experimentally observed kymographs for our two opposing acid–base reservoirs (Fig. 4A), using diffusion coefficients *D*_*a*_ = 9 × 10^−6^ cm^2^/s and *D*_*b*_ = 5 × 10^−6^ cm^2^/s, approximately one order of magnitude lower than their experimental values. We choose the intrinsic reaction rate of acid-base neutralization to be *k* = 104 cm3/mol/s, low enough to prevent numerical instabilities (*SI Appendix*).

To validate that these predicted neutral regions govern network reorganization, we examined how *Physarum* responds within such pH landscapes. Networks actively avoid unfavorable pH regions, leaving little mass in unfavorable pH regions as veins retract and redistribute toward the neutral pH region. To test whether these environments also guide *de novo* network formation from dispersed units, we introduced droplet-like cells into the same pH landscape (Fig. S6). While droplet-like cells fail to persist when exposed to a single acid source (Fig. S3), complex pH landscapes allow them to rapidly fuse and extend protrusions within the neutral zone, forming interconnected networks. In the absence of pH gradients, protrusions form with no obviously preferred direction, whereas in opposing gradients, droplet-like cells consistently adopt elongated morphologies aligned with the predicted neutral region, mirroring the behavior of pre-formed networks.

Introducing multiple fluidic sources of acid and basic solutions enabled the generation of more complex pH landscapes. Reservoirs were carved in grid-like patterns (3 × 3) and filled with pH 2 and pH 12 solutions, while droplet-like cells of *Physarum* microplasmodia were placed within the same configuration (Fig. 4D, Movie S6). Diffusion from these pH reservoirs rapidly established well-defined pH patterns across the agar. Consistent with previously observed directed network formation (Fig. S5), new protrusions emerged at nearneutral zones between unfavorable pH regions. *Physarum* networks expanded preferentially along these favorable pH corridors, and individual droplet-like cells frequently fused upon contact, forming larger interconnected networks. As the pH reservoirs became depleted, pH gradients gradually blurred due to diffusion and neutralization, restoring a nearly homogeneous environment. The imposed pH geometry thus acts as a transient spatial cue that is naturally erased over time. Correspondingly, once gradients dissipate, the initially constrained network structures broaden. While additional geometries (e.g., circular or cross-like patterns) can be generated (Fig. S7, Movies S7, S8), this approach requires physically modifying the agar and injecting fluids, which limits control and reproducibility. To overcome these constraints, we therefore extended our experimental setup.

### Quantitative evolution of diffusive pH through electrochemical control

Using water electrolysis to generate H_3_O^+^ and OH^−^ ions, we created dynamic pH landscapes. We built a 3D-printed mold with a central chamber filled with agar mixed with phenol red, and four connected holes, filled with distilled water and phenol red, see Fig. 5A. Each connected hole hosts a platinum electrode, which can be paired with another electrode. We applied a constant 5 V voltage difference across the electrodes, continuously generating hydroxide ions (OH^−^) at the cathode and protons (H_3_O^+^) at the anode. The increase in ionic concentrations induces an electro-diffusive flux driven by both concentration gradients and an externally applied electric field, which can be described by the Nernst-Planck equation. In our system, the evolution of acid and base concentrations reads:

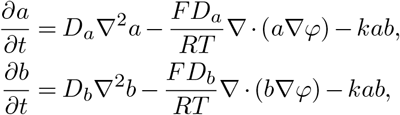

where *F* = 96485 C/mol is the Faraday constant, *R* the perfect gas constant, *T* the room temperature (25°C), and *φ*(*x, y*) is the externally imposed electric potential. This formulation corresponds to an advection-diffusion system with an additional reaction term. From this equation, one can define an effective drift velocity for the ionic species due to the electric field: 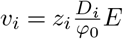, where *i* = *a, b, z*_*i*_ is the ionic valence, *D*_*i*_ the diffusion coefficient, and *E* = Δ*φ/L* the applied electric field over a chamber of length *L* = 20 mm. We further simulate this electro-diffusive system, reproduc-ing qualitatively the observed evolution of the neutral region (Fig. 5B). The velocity of the diffusion fronts follows a linear dependence on the applied voltage, *v*_*i*_ = *α*_*i*_*E*, where *α*_*i*_ = *D*_*i*_/(*φ*_0_ *L*), with *D*_*i*_ the diffusion coefficient, *φ*_0_ the thermal voltage, and *L* = 15 mm the system size. From fits to the experimental data, we obtain *α*_*a*_ = 7.97 × 10^−2^ mm/min/V for the acid front and *α*_*b*_ = 5.23 ×10−2 mm/min/V for the basic front (see Fig. S8).

To experimentally control pH pattern formation, we sequentially activate pairs of electrodes. First, a vertical configuration is applied, with the cathode at the top and the anode at the bottom. This rapidly increases acid and base concentrations in the reservoirs, and within approximately one hour, diffusion into the chamber leads to the formation of a horizontal neutral region (Fig. 5C, Movie S9). After 180 min, this electrode pair is switched off, and a horizontal (left–right) configuration is activated, with the cathode on the left. While diffusion from the top and bottom boundaries persists because of ion accumulation in the reservoirs, the newly imposed field drives ion transport from the left and right boundaries, progressively reorienting the neutral region from horizontal to vertical.

Finally, we use the electrode-based setup to dynamically control the network of *Physarum*. Droplet-like *Physarum* cells were first deposited on the chamber agar, and a DC voltage of 3 V was applied across the horizontal electrode pair (Fig. 5D, Movie S10). The resulting electro-diffusive fronts reached the organisms after ∼270 min and established a neutral zone along which protrusions grew. After 300 min, the horizontal field was switched off, and a vertical electrode pair was activated. This reoriented the pH landscape and induced a corresponding reorganization of the network, with growth fronts relocating to the newly formed neutral zone. The organism remained confined to this dynamically imposed neutral zone for several hours, continuously adapting its morphology to the evolving environment.

Together, these results demonstrate that externally programmable pH landscapes can guide both the formation and reconfiguration of living transport networks. By coupling electro-diffusive control with network redistribution following escape mechanisms, the system achieves continuous, feedback-free tracking of dynamic environments. This establishes a simple framework for external control of living matter, where environmental fields encode spatiotemporal in-structions that are directly interpreted by a vein network.

## Discussion

In this work, we show that distributed transport networks enable efficient detection and response to spatially extended aversive stimuli through the coupling of sensing, transport, and mechanical adaptation. Using *Physarum polycephalum* as a model system, we demonstrate that a networked architecture not only enhances detection of unfavorable conditions but also enables coordinated escape via directed mass redistribution. This response emerges from local mechanical feedback, where contractions in veins exposed to unfavorable conditions generate pressure gradients that drive transport toward favorable areas, resulting in system-level reorganization without centralized control.

By using pH as a tunable external stimulus, we further establish that environmental fields can be precisely structured and dynamically controlled through both diffusion and electrochemical approaches. This enables the generation of complex, time-dependent pH landscapes that guide network architecture and function. Together, these results highlight how distributed architectures can integrate spatial information and convert it into adaptive behavior, providing a physical basis for embodied sensing and control in networked systems.

### Limitations of the study

While *Physarum* provides a powerful and accessible model for studying distributed sensing and adaptation, several limitations must be considered when generalizing these findings.

First, the scale of the system is enabled by the unicellular yet multinucleated nature of *Physarum*, allowing transport and coordination over macroscopic distances. In contrast, most biological systems operating at comparable scales are multicellular and rely on communication between discrete cells. Such distributed coordination mechanisms are conceptually closer to swarm robotic systems, where large populations of individual agents collectively achieve robustness, self-organization, and adaptive behavior without centralized control (44, 45). Swarm robotic systems can furthermore exhibit morphogenetic behaviors and collective restructuring through local interactions between agents (46). *Physarum* fundamentally differs from these systems in that coordination emerges not from communication between discrete units, but through continuous transport, mechanical coupling, and flow-mediated signaling within a single deformable body. Rather than comprising many agents, the organism realizes a distributed network of sensor-actuator elements whose number, connectivity, and spatial arrangement evolve over time. This enables a single agent to exhibit behaviors commonly associated with multi-agent systems, including decentralized coordination, adaptive reconfiguration, and robustness to local perturbations. In this sense, *Physarum* occupies an intermediate position between soft robotic materials and swarm robotic systems, suggesting a possible route toward soft robotic architectures that combine continuous embodied mechanics with swarm-like distributed adaptation.

Second, the pH ranges explored in this study (pH 2-12) represent extreme conditions that are toxic for most biological systems, although, e.g., insect-based biohybrid systems have shown remarkable resilience to extreme pH environments (47). While these pH conditions provide a robust and well-controlled stimulus to *Physarum*, they limit direct applicability to other systems. However, electrochemical approaches offer a route toward finer control, where local gradients could be generated using microelectrodes and reduced current intensities to create biologically and technologically relevant conditions. Combined with other stimuli such as light (48) or nutrients (36), pH control may offer a versatile platform to guide and test adaptive behavior.

In addition, our experiments are performed in quasi-two-dimensional environments, whereas many real-world applications involve fully three-dimensional networks. Extending these principles to 3D architectures remains an important and likely direction, particularly for applications in soft robotics and tissue engineering, as the underlying transport mechanisms are not restricted to two-dimensional geometries.

Finally, while we identify a physical mechanism linking local vein contraction to global transport and adaptation, we did not study the underlying molecular pathways governing stimulus detection and response. Response to positive stimulus, such as food, was proposed to be mediated by a softening agent, hypothesized to be calcium ion Ca^2+^, that triggered a response of tube dilation near the stimulus (36, 49). Although not required to establish the mechanical principles described here, such insights could facilitate the translation of these behaviors into metamaterial or synthetic biohybrid systems with tunable response characteristics (50, 51). Promising results have already been achieved with responsive hydrogels, especially pH-sensitive hydrogels, which can produce continuous vein contractions when exposed to acidic conditions (52, 53). More broadly, *Physarum* remains a living system whose internal processes can hardly be directly programmed at the microscopic level, highlighting the need for engineered analogs that retain similar functionality while offering greater controllability. These aspects point toward several promising directions at the interface of soft robotics, biohybrid systems, and active materials.

### Outlook and impact

One immediate avenue of our work is the development of synthetic transport networks that replicate the coupling between flow, structure, and environmental sensing observed here. Advances in fluidic soft robotics, hydrogel systems, and electroactive materials suggest that such architectures could be engineered to achieve distributed sensing and adaptive reconfiguration (54, 55). In this context, environmental fields, chemical, electrical, or mechanical, could serve as programmable inputs that dynamically guide system behavior without centralized control. Chemical gradients are already used to direct microscale robotic systems; for example, pH taxis has been employed to position and guide biohybrid microrobots in chemically structured environments (56). Our results extend this concept from the control of discrete agents to the regulation of continuously reconfigurable transport networks.

Beyond fully synthetic systems, biohybrid approaches offer an alternative pathway. Embedding living transport networks within engineered platforms could enable systems that sense and repair damage (57), redistribute resources, and reconfigure autonomously, similar to *Physarum* (41). For instance, vascularized robotic materials or living circuits incorporating *Physarum*-like networks could combine transport, sensing, and computation within a single substrate, enabling closed-loop interactions with electronic components and adaptive responses to environmental changes.

More broadly, our results suggest that distributed transport networks can serve as a platform for embodied computation, where information processing is not localized but emerges from physical interactions between flow, structure, and environment (58, 59). This perspective opens opportunities for designing systems in which control is encoded in material organization and external fields, rather than in discrete algorithms.

Finally, *Physarum* itself may serve as a testbed for exploring such concepts. Its accessibility, scalability, self-organized hierarchy, and dynamic network behavior provide a unique platform to investigate how particles, microrobots, or chemical agents can be transported, distributed, and coordinated within a living flow network. Such systems could inform strategies for targeted delivery, adaptive routing, or collective behavior in complex environments.

Taken together, these findings point toward a new class of autonomous systems in which sensing, computation, and actuation are intrinsically coupled through distributed transport. By leveraging environmental fields as control signals and exploiting the physical principles of network adaptation, it may be possible to design soft robotic systems capable of robust operation in dynamic and unstructured environments.

## Materials and Methods

### Preparation of pH solutions

Solutions with defined pH values were prepared by adjusting the pH of a 100 mM HEPES buffer (PUFFERAN, Carl Roth) using an electrode-based pH meter (Mettler Toledo FiveGo F2). For acidic solutions, hydrogen chloride (Sigma Aldrich 7647-01-0) was added, and for basic solutions, potassium hydroxide (VWR CAS 1310-58-3) was used. To ensure the pH stability of the solutions and to visually characterize diffusion kinetics, dyes such as Phenol Red (ThermoScientific CAS 143-74-8), Thymol Blue (ThermoScientific CAS 76-61-9), Bromophenol Blue (ThermoScientific, CAS 115-39-9), Bromothymol Blue (ThermoScientific, CAS 76-59-5) and Methyl Red (Thermo-Scientific, 493-52-7) were added.

### Cultivation of *Physarum polycephalum*

*Physarum polycephalum* was cultivated from microplasmodia extracted from sclerotia (Carolina Biological Supply Company) under axenic conditions, as detailed in (60). Briefly, microplasmodia were incubated at 25°C in liquid medium composed of Balanced Salt Solution and Semi-Defined Medium, contained in shaking Erlenmeyer flasks. A new passage was created every two days by transferring 1 mL of the previous culture into 100 mL of fresh medium. Prior to plating, microplasmodia were centrifuged in an Eppendorf tube for 30 seconds, and the supernatant was removed. Up to eight drops (approximately 50 *µ*L each) of the remaining microplasmodial content were placed onto 2% agar (VWR Agar for bacteriology 9002-18-0) supplemented with 15 mg L^−1^ Phenol Red unless otherwise specified. The plated microplas-modia were incubated at 25°C for approximately 15 hours, allowing network formation. For patterning experiments, the resulting networks were cut into a square shape using a plastic inoculation loop (VWR).

### Microscopy imaging

Plasmodia were imaged with a Zeiss Axio Zoom V.16 microscope equipped with a Hamamatsu ORCA-Flash 4.0 digital camera for grayscale imaging, AxioCam305 for color imaging, and a Zeiss PlanNeoFluar 0.5x objective. The Zeiss Zen 3.7 (Pro) software was used for imaging.

### pH patterning setup

pH patterns used to guide the network shape of *Physarum polycephalum* were generated in standard Petri dishes filled with 2% agar. Reservoirs for pH solutions were created by carving holes into the agar using a biopsy punch, cutting straight shapes with a scalpel, or pressing 3D-printed PLA templates into the gel. The templates were fabricated using a Bambulab X1E printer. Reservoirs were filled with the pH solutions prepared as described above, which then diffused into the agar gel. To ensure solution stability, all pH solutions contained HEPES buffer, while the agar itself was prepared without buffer to allow controlled adjustment of the local pH over relevant timescales.

### Electrolytic setup

Electrolysis was conducted using Platinum electrodes (Sigma Aldrich, 7440-06-4, 0.5 mm), soldered with tin to electric wires. To provide electric current, we used two power generators (VoltCraft DSP-3005, RIGOL DG812).

## Data availability

Experimental datasets and scripts used to perform data analysis are stored on MediaTUM under https://doi.org/10.14459/2026mp1858055, an open-access repository provided by the Technical University of Munich.

### Use of AI-assisted technologies

During the preparation of this work, ChatGPT (OpenAI; GPT-5) was used to assist in writing portions of the code for tracking and segmenting droplet-like cells. All AI-assisted code was reviewed, tested, and validated by the authors before use. Grammarly (Grammarly Inc.) was used to check grammar and language in the manuscript text; the authors reviewed and take full responsibility for the final text.

## ACKNOWLEDGEMENTS

We thank David Zwicker for his invaluable help on using the Py-PDE package, and the BPM and E14 team members for their support and discussions. Funded by the German Federal Ministry of Research, Technology and Space (BMFTR) in the program of “Souverän. Digital. Vernetzt.” (to F.C.S.). Joint project 6G-life, project identification number 16KISK002 (to F.C.S.); the European Research Council (ERC) under the European Union’s Horizon 2020 research and innovation program (grant agreement No. 947630, FlowMem) (to K.A.); the Deutsche Forschungsgemeinschaft DFG through INST 95/1634-1 FUGG (to K.A.); and the Deutsche Forschungsgemeinschaft (DFG, German Research Foundation) under Germany’s Excellence Strategy – EXC3092/1 – 533751719.

## Authors contribution

Conceptualization: M.L.V.S., A.B., F.C.S., K.A. Methodology: M.L.V.S., A.B., F.C.S., K.A. In-vestigation: M.L.V.S., A.B. Visualization: M.L.V.S., A.B. Funding acquisition: F.C.S., K.A. Project administration: F.C.S., K.A. Supervision: F.C.S., K.A. Writing – original draft: M.L.V.S., A.B., F.C.S., K.A. Writing – review & edit-ing: M.L.V.S., A.B., F.C.S., K.A.

M.L.V.S. and A.B. contributed equally to this work.

### Authors declaration

The authors declare that they have no competing interests.

