## Supplementary Information for "Distributed network sensing as an escape strategy"

[a]Technical University of Munich, School of Natural Sciences
[†]These authors contributed equally to this work

**Reaction–diffusion model: physical parameters and time scales.** We model the interaction between an acid and a base using reaction–diffusion equations:

$$\frac{\partial a}{\partial t} = D_a \nabla^2 a - kab, \tag{S1}$$

$$\frac{\partial b}{\partial t} = D_b \nabla^2 b - kab, \tag{S2}$$

where $a(x,y,t)$ is the acid concentration, $b(x,y,t)$ is the base concentration, $D_a$ and $D_b$ are the diffusion coefficients of acid and base, and $k$ is the second-order reaction rate constant. All quantities are expressed in centimetre-based units for consistency. For the left/right diffusion setup, we impose Neumann boundary conditions, such that $(\partial a/\partial x)_{x=0} = J_a/D_a$ (positive flux of acid on the left), $(\partial b/\partial x)_{x=L} = J_b/D_b$ (positive flux of base on the right), and zero (reflective) otherwise. We assume $J_a = J_b$ for symmetry and mass conservation.

**Physical parameters.** The domain size corresponds to the physical size of the experimental chamber and sets the characteristic diffusion length scale. Typically, we use $L = 15$ mm. The diffusion coefficients for the ions diffusing in water are typically small for acid and base, of the order of $D_{a,b} \sim 10^{-4}\,\mathrm{cm}^2/\mathrm{s}$. For the reference concentration $c_0$, we convert the pH values in $\mathrm{mol/cm}^3$, as $[\mathrm{H_3O^+}] = 10^{-\mathrm{pH}} = 10^{-3}\,\mathrm{mol/cm}^3$. We obtain, for pH = 2, a concentration of $10^{-5}\,\mathrm{mol/cm}^3$, which represents the maximum concentration of the system. Thus, we choose the reference concentration $c_0 = 10^{-4}\,\mathrm{mol/cm}^3$. Finally, the reaction rate $k$ is reported with extremely high values, $k = 10^{14}\,\mathrm{cm}^3/\mathrm{mol/s}$.

**Characteristic time scales.** Two competing timescales govern the system dynamics.
(i) The diffusion time, $\tau_D = L^2/D$, required for diffusion to transport species across the system, typically $\tau_D \sim 10^4\,\mathrm{s} = 160$ min at our length scales for hydroxide ($\mathrm{OH^-}$) and protons ($\mathrm{H^+}$);
(ii) the reaction time, $\tau_R = 1/(kc_0)$, the time required for neutralization at concentration $c_0$, typically $\tau_R \sim 10^{-10}\,\mathrm{s}$, almost instantaneous. Thus, we find a **Damköhler number**, which measures the relative importance of reaction and diffusion:

$$Da = \frac{\tau_D}{\tau_R} = \frac{kc_0 L^2}{D} \sim 10^{15}, \tag{S3}$$

to be extremely large, therefore indicating the formation of a sharp front. However, to avoid numerical instabilities, we reduce the value of the reaction rate to $k = 10^4\,\mathrm{cm}^3/\mathrm{mol/s}$, which gives a reaction time $\tau_R = 1$ s, still lower than the diffusion timescale.

**Numerical implementation.** To integrate the PDEs, we use py-pde, a Python library specifically created to solve the time-evolution of spatial PDEs through the finite-difference method, with several boundary conditions. py-pde allows one to generate simple geometries, and to define equations for the PDE with symbolic notations. In detail, we typically generate a 2D Cartesian grid, with size $(N_x \times N_y)$. The concentrations $a(x,y,t)$ and $b(x,y,t)$ are represented as scalar fields in a field collection. Boundary conditions are then defined, either Dirichlet, Neumann or Robin. We typically use Neumann boundary conditions, such that there is a given flux of concentration to form pH patterns. The library allows for more complex boundary conditions, such as space- or time-dependent functions. We then define the system of PDEs to match Eqs. Eq. (S1)–Eq. (S2). The system can then be integrated using the built-in solver of the library, using adaptive time-stepping, and recording the concentrations at each pixel in a dedicated memory environment. We do not rescale the equations unless specified, to keep actual units for the solution and a direct mapping to the experiments.

**Annihilation process and scaling of the neutral zone.** The system of PDEs, Eqs. Eq. (S1)–Eq. (S2), corresponds to a two-species annihilation system (1). Since our setup is mostly one-dimensional (because of the imposed boundary conditions), we only study the $x$-direction. The system starts as neutral at first, i.e., $a, b \sim c_0$. As time proceeds, diffusion fronts develop as circles, and their dynamics can be described with diffusion only, since the two species do not meet yet. Eventually, they meet roughly in the centre.

This type of reaction-diffusion system leads, in its steady state, to a characteristic profile of the concentrations. One can define three regions: on the left (resp. on the right), a freely diffusing region of species A (resp. species B), with a linear profile of concentration, and in the centre, a quick decay where the species meet.

In steady state, the left- and right-most regions are mostly controlled by pure diffusion of only one species since only one species is present: $kab \ll D_a \nabla^2 a$, $D_b \nabla^2 b$, which leads to $a(x) \sim -(J_a/D_a)x + a_0$ and $b(x) \sim (J_b/D_b)x + b_L$, where $a_0$ and $b_L$ are constants determined by the boundaries. They need to be equal, $a_0 = b_L$, as there need to be as many particles of $a$ as of $b$ entering the system to have a steady state.

The reaction region, defined as $R(x) = ka(x)b(x)$, is progressively created, evolving from noisy initial conditions to bimodal distributions meeting at the front, and later forming a Gaussian distribution, see Fig. S1A.

At steady state, the flux of ions balances, and concentrations of both species are non-negligible, and should be of order $a, b \sim (J/D)w$, where $w$ is the width of the reaction region. To get a scaling of $w$, we calculate the number of annihilation events by integrating the reaction term, $R(x)$, over the region:

$$N_{\text{annihilations}} = \int_{-w/2}^{w/2} R(x)\,dx \sim kabw \sim J, \tag{S4}$$

since it should be compensated by the number of particles entering the domain. This leads to the scaling for the width of the reaction region

$$w \sim \left(\frac{D_a D_b}{Jk}\right)^{1/3}. \tag{S5}$$

Below, we show the scaling of $w$ for several conditions of the reaction rate $k$, see Fig. S1B. We notice that for small rates, $k < 10^2\,\mathrm{cm}^3/\mathrm{mol/s}$, the scaling fails, which is expected since there is not much interaction between the species. Further, there exists a lower bound on the flux of the system: the length $w$ cannot exceed the system length $L$, and since $k$ and $D_a$, $D_b$ are fixed, the bound is on the flux $J$, and obeys

$$J > J_0 = \frac{D_a D_b}{kL^3}, \tag{S6}$$

and we also observe such an effect in the simulations, see Fig. S1C.

### Electrochemical kinetics.

***Anode & cathode reactions..*** In our setup, we use an electrolytic cell, that requires electric current to drive chemical reactions. The anode and cathode are, in our case, made of platinum, which means there is no corrosion of the electrodes – unlike Ag-AgCl electrodes. The cathode is where the reduction happens, and obeys the chemical reaction

$$2\mathrm{H_2O}_{(l)} + 2e^- \rightarrow \mathrm{H}_{2(\mathrm{g})} + 2\mathrm{OH}^-_{(aq)}, \tag{S7}$$

therefore, the cathode produces alkaline solution and hydrogen gas. The anode is where the oxidation happens, and obeys the chemical reaction

$$2\mathrm{H_2O}_{(l)} \rightarrow \mathrm{O}_{2(\mathrm{g})} + 4\mathrm{H}^+_{(aq)} + 4e^-, \tag{S8}$$

therefore, the anode produces acid solution and oxygen gas.

***Nernst–Planck–Poisson equations..*** Compared to our previous reaction-diffusion system, the introduction of an electric voltage source induces coupling between diffusion and electrostatics, and introduces a drift term for the ions due to the electric field $E = -\nabla\varphi$, in the Fick flux: $J_i = -D_i\left(\nabla c_i + \frac{z_i F}{RT} c_i \nabla\varphi\right)$, where $F = 96485$ C/mol is Faraday's constant, $c_i = a, b$ is the concentration of ions, and $z_i = \pm 1$ is the valence of each ion. Our previous PDEs for the evolution of $a$ and $b$ become:

$$\frac{\partial a}{\partial t} = D_a \nabla^2 a + \frac{D_a F}{RT} \nabla \cdot (a \nabla \varphi) - kab, \tag{S9}$$

$$\frac{\partial b}{\partial t} = D_b \nabla^2 b + \frac{D_b F}{RT} \nabla \cdot (b \nabla \varphi) - kab, \tag{S10}$$

$$\nabla^2 \varphi = \frac{F}{\varepsilon}(a - b), \tag{S11}$$

where $\varepsilon$ is the permittivity of water and the last equation is the Poisson equation, which describes the coupling and the influence of the ions on the potential $\varphi$. Note that non-dimensionalizing the Poisson equation, using $\bar{x} = x/L$, $\bar{a} = a/c_0$ and $\bar{\varphi} = \varphi/\varphi_0$

where $\varphi_0 = RT/F \simeq 25$ mV, we extract the Debye length $\lambda_D^2 = RT\varepsilon/(F^2 c_0)$, and the Poisson equation becomes

$$\nabla^2 \bar{\varphi} = \frac{L^2}{\lambda_D^2}(\bar{a} - \bar{b}). \tag{S12}$$

Using $\varepsilon = 710 \times 10^{-12}\,\mathrm{C/(V\,m)}$ for the permittivity of water, $T = 298.15$ K (at 25°C), $c_0 = 10^2\,\mathrm{mol/m^3}$, we obtain $\lambda_D = 1.89$ nm $\ll L$. This indicates that the electric boundary layer is extremely thin and that most of the system is in the bulk, where electroneutrality prevails, and any charge imbalance creates a large potential curvature. Therefore, the Poisson equation reduces to the Laplace equation $\nabla^2 \varphi = 0$, which gives a linear profile for the electric potential, simplifying our system of PDEs. Assuming the electric potential is fixed, one can find the drift velocity for each ion as $v_i = -z_i \frac{D_i F}{RT} E$, and the PDEs become

$$\frac{\partial a}{\partial t} = D_a \nabla^2 a + v_a \nabla a - kab, \tag{S13}$$

$$\frac{\partial b}{\partial t} = D_b \nabla^2 b + v_b \nabla b - kab. \tag{S14}$$

**Tube versus fork resistance.** We consider two graphs, as depicted in Fig. 3D of the main text: both have three edges, depicted in black, but differ in topology. These graphs represent flow networks, where the edges are cylinders that can transport a laminar fluid flow using pressure differences.

We refer to the two graphs as tube (T) and fork (Y). Each edge is associated with a hydraulic resistance $R$. As boundary conditions, we introduce two more nodes, with fixed pressures, acting as sources ($p^+$) and sinks ($p^-$). The sinks and sources are connected to the graph with edges that have a different resistance, $R^+$ and $R^-$. To calculate the efficiency of each network, we compare the net flow $\Phi_{T,Y}$ that exits the system through the sinks.

From there, we write Kirchhoff's law for each of the nodes, i.e., flux conservation: for a node $i$, it reads

$$\sum_j Q_{ij} = 0. \tag{S15}$$

Using Ohm's law, $\Delta P_{ij} = R_{ij} Q_{ij}$, where $Q_{ij}$ is the flow in tube $(i,j)$, $\Delta P_{ij}$ is the pressure drop from node $i$ to node $j$ and $R_{ij}$ is the hydraulic resistance, and after some algebraic calculations, one can express the net flow $\Phi_{T,Y}$ as

$$\Phi_T = \frac{p^+ - p^-}{R^+ + R^- + 3R}, \tag{S16}$$

$$\Phi_Y = \frac{2(p^+ - p^-)}{R^+ + 2R^- + 3R}. \tag{S17}$$

The ratio

$$\frac{\Phi_T}{\Phi_Y} = \frac{R^+ + 2R^- + 3R}{2(R^+ + R^- + 3R)} < 1 \tag{S18}$$

indicates that for similar hydraulic resistances, the fork topology Y will evacuate more flow than the tube topology T. If $R^+ = R^- = R$, the ratio is 0.6.

**References.**


1. E. Ben-Naim and S. Redner. Inhomogeneous two-species annihilation in the steady state. *J. Phys. A Math. Gen.*, 25:L575–L583, 1992.

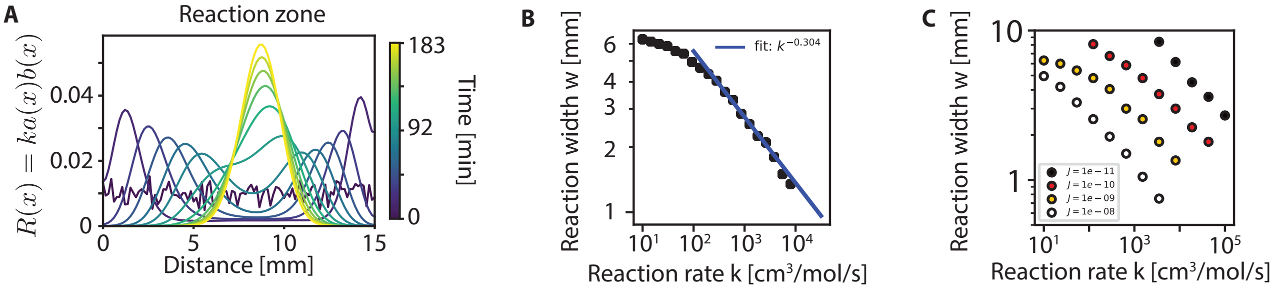


**Fig. S1. Reaction region's behaviour.** (**A**) Reaction zone evolution for several time points shows a convergence towards a Gaussian. (**B**) Reaction zone half-width versus reaction rates with a fit. (**C**) Reaction zone half-width versus reaction rates for several fluxes.

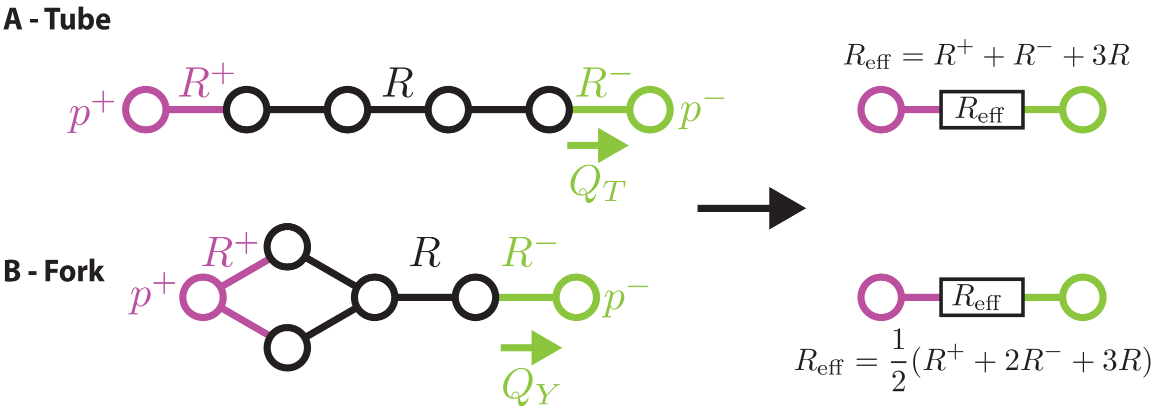


**Fig. S2. Tube versus fork architectures.** (**A**) Tube versus (**B**) fork architectures, with fixed pressures at the boundaries. Purple represents sources, green represents sinks.

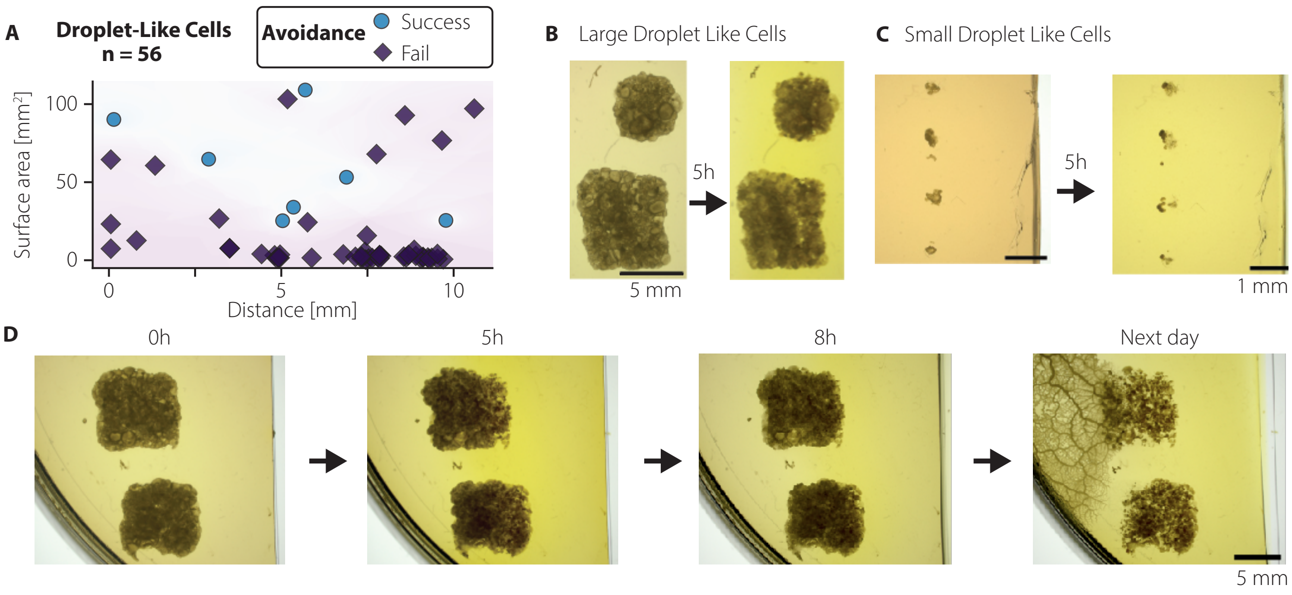


**Fig. S3. Droplet-like cells fail to respond to harmful pH gradient.** (**A**) Success or failure to avoid the advancing aversive pH 2 gradient for (**B**) large droplet-like cells (scale bar 5 mm) and (**C**) small droplet-like cells (scale bar 1 mm). (**D**) Successful escape of a large droplet-like cell from acid after 24 hours, by forming a network.

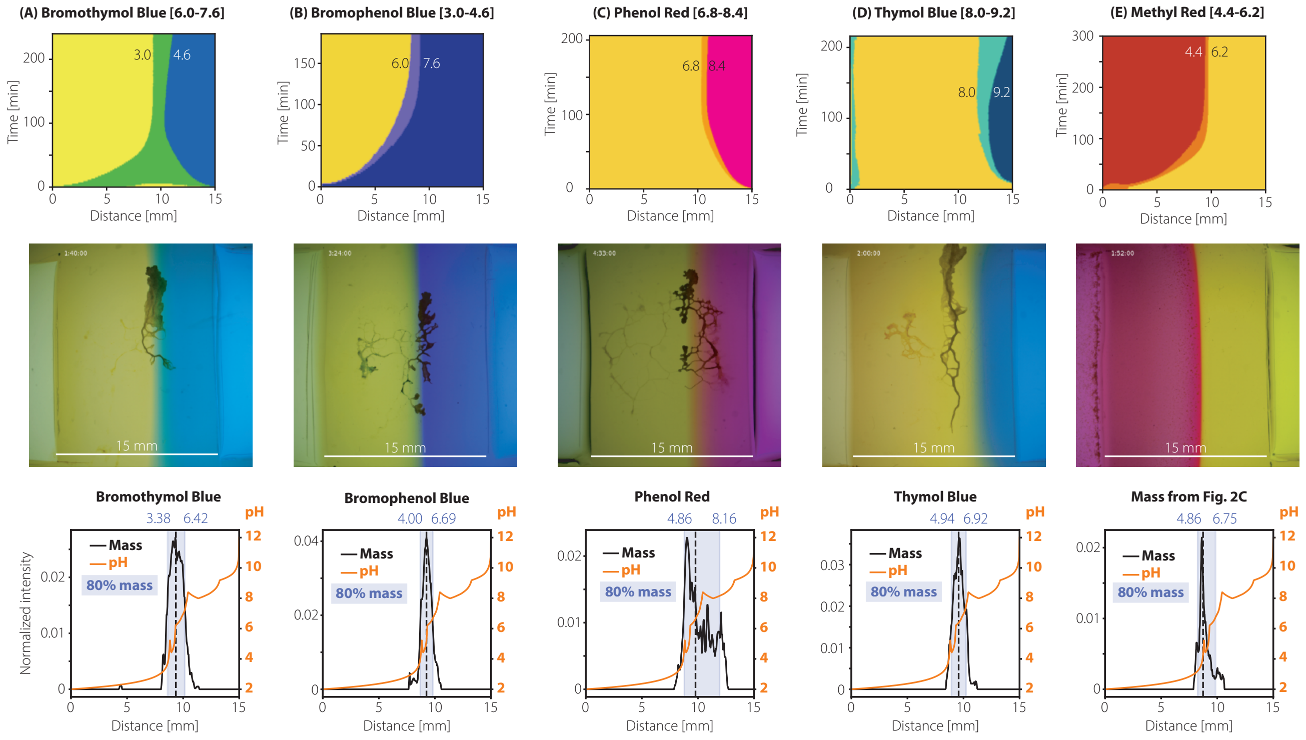


**Fig. S4. Extracted kymographs of pH dyes and corresponding microscope images with *Physarum* networks on top.** Dyes are (with transition ranges): (**A**) Bromophenol Blue (3.0-4.6); (**B**) Bromothymol Blue (6.0-7.6); (**C**) Phenol Red (6.8-8.4); (**D**) Thymol Blue (8.0-9.2); (**E**) Methyl Red (4.4-6.2) was not used with *Physarum* given its cytotoxicity and the precipitates that formed. On the middle line, we show the *Physarum* networks used to calculate the mass distribution of the bottom line. On the bottom line, we plot the mass distribution of *Physarum* vs. distance, with the reconstituted pH profile. Extracting 80% of the mass, we find preferred regions for *Physarum* in each condition.

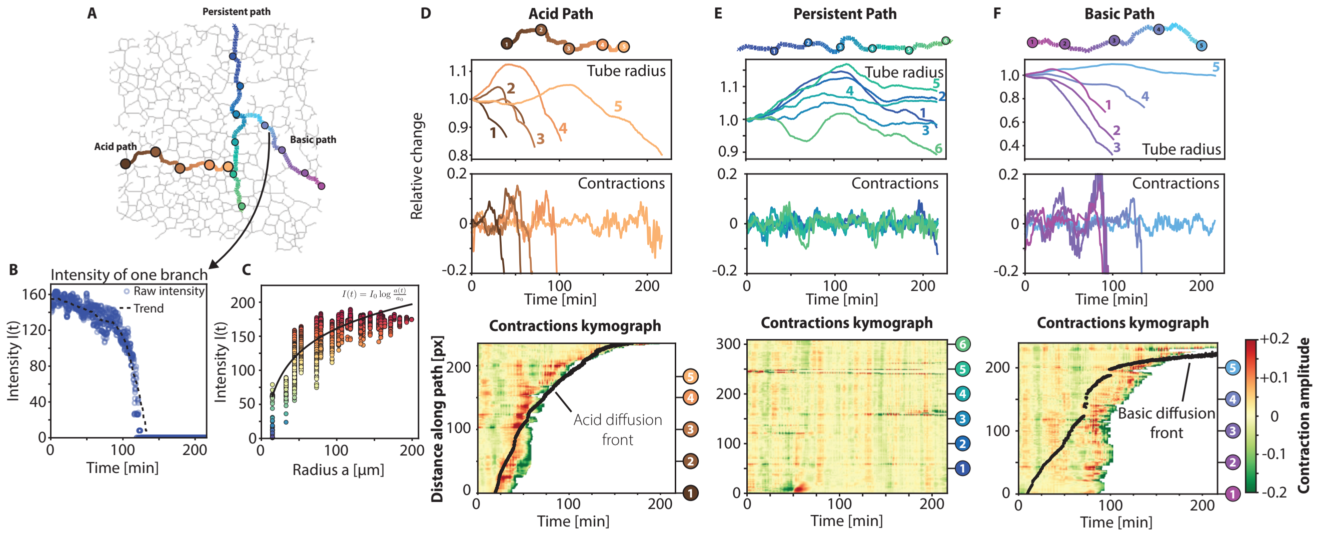


**Fig. S5. Contraction dynamics show a response to acid and basic stimuli.** (**A**) Defining paths spanning three directions (acid, basic and persistent) along the network skeleton, (**B**) we extract the pixel intensity over time, here plotted for a representative point in the basic path. The trend reflects the averaged signal, and (**C**) can be mapped to the local radius using the Beer-Lambert law. (**D–F**) For each of the paths, we extract the trend (averaging over 20 min) and contractions of vein radii at specific locations along the paths and show the kymographs of contractions along the entire paths with the respective diffusion front superimposed. While the acid path shows an increase in contractions before the acid wave arrives, the basic path is delayed in its response. The persistent path remains unperturbed.

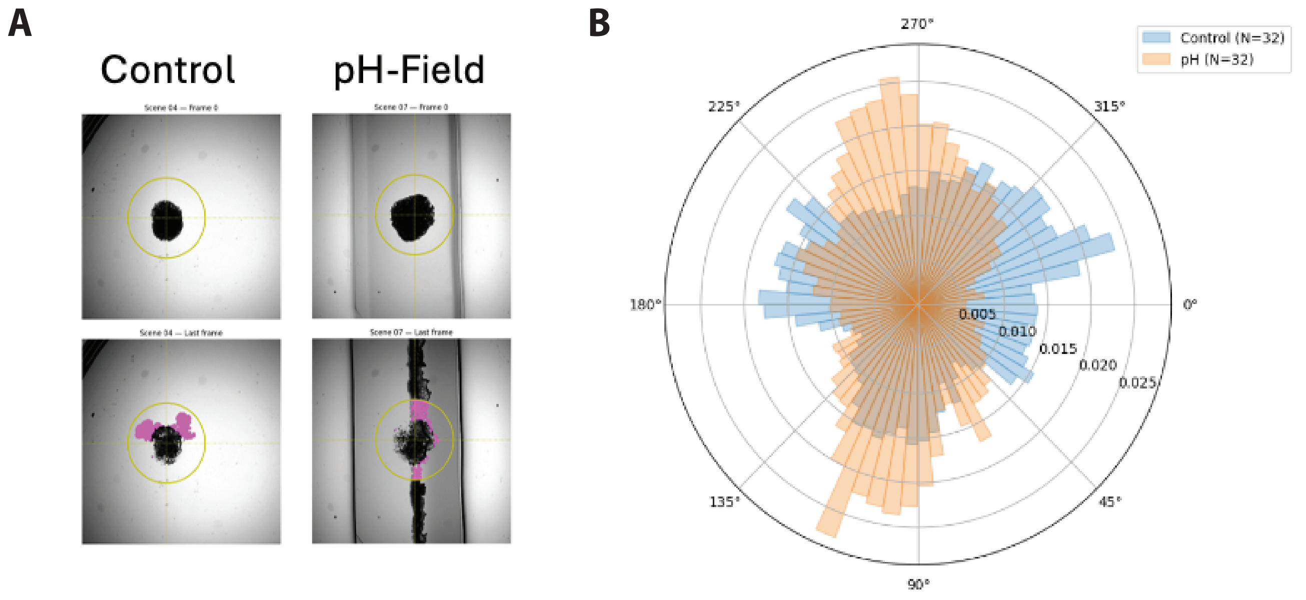


**Fig. S6.** ***De novo Physarum* networks preferentially form protrusions in the neutral regions.** (**A**) (Left) Control droplet-like cells in a pH-free environment extend protrusions in all directions, while (Right) droplet-like cells form protrusions and generate a network along the neutral region only when exposed to acid & base stimuli. (**B**) Quantification of the protrusion orientation in opposing pH gradients is vertical to the pH gradient axis.

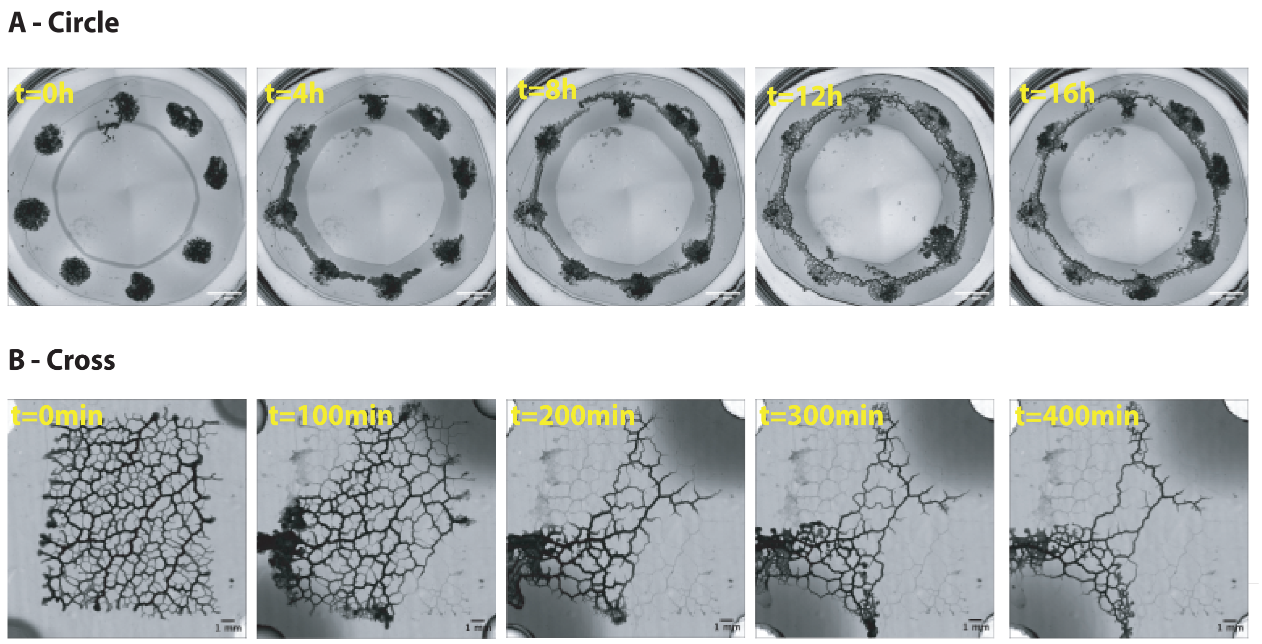


**Fig. S7. Fluidic control of vein network architecture.** We showcase on-demand network architecture, exemplified by a (**A**) circle network and (**B**) a cross network, obtained by carving reservoirs and filling them with acid (pH = 2) and alkaline (pH = 12) solutions.

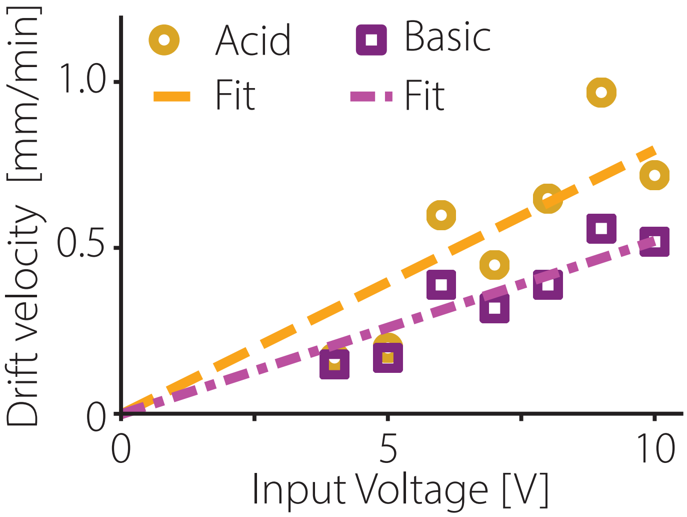


**Fig. S8. Effective drift velocity of acid and basic fronts versus input voltage**, in volts, for acid and basic fronts.

### Supplementary Movies

**Movie S1.** *Physarum* network on agar + Phenol Red substrate. Stimuli are pH 2 (left) and pH 12 (right), contractions are visible. Frame rate 10 s. Left-right width 15 mm.

**Movie S2.** *Physarum* network on agar + Bromothymol Blue substrate. Stimuli are pH 2 (left) and pH 12 (right). Frame rate 2 min.

**Movie S3.** *Physarum* network on agar + Bromophenol Blue substrate. Stimuli are pH 2 (left) and pH 12 (right). Frame rate 2 min.

**Movie S4.** *Physarum* network on agar + Phenol Red substrate. Stimuli are pH 2 (left) and pH 12 (right). Frame rate 1 min.

**Movie S5.** *Physarum* network on agar + Thymol Blue substrate. Stimuli are pH 2 (left) and pH 12 (right). Frame rate 2 min.

**Movie S6.** *Physarum* droplet-like cells on agar + Phenol Red substrate, forming a grid-like pattern. Wells are filled with pH 2 and pH 12. Frame rate 3 min.

**Movie S7.** *Physarum* droplet-like cells on agar + Phenol Red substrate, forming a circle-like pattern. Greyscale camera is used. Wells are filled with pH 2 (outside ring) and pH 12 (inside ring). Frame rate 3 min.

**Movie S8.** *Physarum* network on agar + Phenol Red substrate, forming a cross-like pattern. Greyscale camera is used. Wells are filled with pH 2 (top-left, bottom-right) and pH 12 (bottom-left, top-right). Frame rate 10 s.

**Movie S9.** Empty 3D chamber. Input voltage is DC 5 V. Top-bottom electrodes are on from $t = 0$ min to 180 min, then left-right from $t = 180$ min to 266 min. Frame rate 10 s.

**Movie S10.** 3D chamber with *Physarum* droplet-like cell and electrodes. Input voltage is DC 3 V. Left-right electrodes are on from $t = 0$ min to 306 min, then bottom-top from $t = 306$ min to 1312 min. Frame rate 1 min.
